# Studying cellular functions in bipolar disorder: Are there specific predictors of lithium response?

**DOI:** 10.1101/753574

**Authors:** Pradip Paul, Shruti Iyer, Ravi Kumar Nadella, Rashmitha Nayak, Anirudh S. Chellappa, Sheetal Ambardar, Reeteka Sud, Salil K. Sukumaran, Meera Purushottam, Sanjeev Jain, ADBS Consortium (ADBS: The Accelerator program for Discovery in Brain disorders using Stem cells), Biju Viswanath

**Affiliations:** The National Institute of Mental Health and Neurosciences (NIMHANS), India; Institute for Stem Cell Biology and Regenerative Medicine (InStem), India; The National Centre for Biological Sciences (NCBS), India; National Institute of Mental Health and Neuro Sciences (NIMHANS), India; National Center for Biological Sciences (NCBS), India

## Abstract

**Background:** *Lithium* is the first-line mood stabilizer for the treatment of bipolar disorder (BD). In order to interrogate cellular phenotypes related to disease and lithium treatment response, this study used neural precursor cells (NPCs) and lymphoblastoid cell lines (LCLs) from BD patients who are well characterized for clinical lithium response.

**Methods:** BD *patients* diagnosed according to the DSM-IV criteria; were recruited from the outpatient services of the National Institute of Mental Health and Neurosciences (NIMHANS), Bangalore, India. Clinical lithium response was assessed using the “Alda scale” and “NIMH Retrospective Life chart method”. The controls were ethnically matched healthy subjects with no family history of neuropsychiatric illness. NPCs from two BD patients from the same family who clearly differed in their clinical response to lithium were chosen, and compared with healthy population controls. Whole transcriptome sequencing (RNA-Seq) and analysis were performed, with and without *in vitro* lithium (1mM for 7 days). In addition, mitochondrial membrane potential (MMP), cell viability and cell proliferation parameters were examined. Experiments were also performed in 25 LCLs from BD patients (16 lithium responders and 9 lithium non-responders), and 12 healthy control LCLs, to evaluate them in a system amenable to clinical translation.

**Results:** RNA-*Sequencing* and analysis did not reveal differences in NPCs on *in vitro* lithium treatment. MMP was lower in BD, both in NPCs and LCLs; reversal with *in vitro* lithium happened only in LCLs and was unrelated to lithium response. Cell proliferation was higher in BD compared to controls, and there was no change on lithium addition. Cell viability assays indicated greater cell death in BD; which could only be rescued in LCLs of clinical lithium responders. The latter finding was associated with enhanced *BCL2* and *GSK3B* expression with *in vitro* lithium.

**Discussion:** *Overall*, our study findings indicate that there are cellular phenotypes related to the disease (mitochondrial potential, cell proliferation) in NPCs and LCLs. We also observed clinical lithium response related phenotypes (cell viability, *BCL2/ GSK3B* expression) in LCLs. The next step would be to evaluate a larger set of PBMCs from clinical lithium response groups of BD to derive cellular phenotypes related to direct clinical application.

## Background

Bipolar disorder (BD) is highly heritable with a lifetime prevalence of 1-3% [1]. It is a neurodevelopmental disorder which manifests in adulthood [2]. While lithium and valproate have been the mainstay for treatment of BD [3] their precise mechanisms of action and pharmacogenomics are poorly understood. Response to lithium varies considerably among patients, and 40-50% patients do not respond to the drug. Factors governing lithium response in BD may be familial [4]. Currently, identification of patients whose symptoms respond favorably to lithium relies upon clinical criteria that lack specificity and sensitivity. In our experience, lithium non-responders tend to be more severely ill [5]. Despite this, lithium remains the ‘gold-standard’ for treatment of BD, and reliably predicting lithium response using molecular markers would be useful.

Most studies have used lymphoblastoid cell lines (LCLs) to study lithium response *in vitro* [6]. However, only a few studies have modelled clinical response to lithium in BD. The primary advantage of identifying molecular markers in lymphocytes/ LCLs would be the potential for clinical translation. However, cells of the neuronal lineage would be most appropriate for studying biological determinants of BD and the mechanisms of lithium action. Human induced pluripotent stem cell (iPSC) models allow for the derivation of specific brain cell types that harbor the patient’s genetic background, thus recapitulating the disease in a physiological context. As reviewed recently [2], studies in iPSC-derived NPCs and neurons have found abnormalities in neural patterning, post-mitotic calcium signaling, and neuronal excitability.

We have used iPSC-derived neural precursor cells (NPCs) of BD patients from families with multiple affected members who differed in their clinical response to lithium and compared these to healthy population controls, to identify cellular phenotypes in relation to BD and response to lithium. These phenotypes were further studied in larger samples of LCLs from BD patients characterized for lithium response. Reversal of phenotypes was attempted with *in vitro* lithium and valproate; the latter being the drug of choice for clinical lithium non-responders in our cohort.

A hypothesis-free approach using RNA-Seq analysis did not reveal genome-wide gene expression differences in NPCs with and without *in vitro* lithium. A hypothesis-based approach based on current literature (supplementary table 1) found cellular phenotypes related to disease [mitochondrial membrane potential (MMP) and cell proliferation] in NPCs and LCLs; and lithium treatment response related phenotypes (cell viability and *BCL2/ GSK3B* expression) in LCLs.

## Materials and Methods

### Clinical recruitment

All BD patients were recruited as part of a study which systematically characterized 210 patients for clinical lithium response [5]. Family A (figure 1) had two BD patients clearly discordant for clinical lithium response (BD1 – non-responder and BD2 – responder), and was recruited as part of a family-based cohort study of psychiatric illness in the Indian population, the Accelerator program for Discovery in Brain disorders using Stem cells (ADBS) [7]. All patients were assessed for clinical lithium response using the Alda Scale and NIMH Retrospective Life chart method [4, 8]. A subset of 25 BD patients who had extreme phenotypes for clinical lithium response [Lithium responders with Alda score ≥7 (N=16) and lithium non-responders with Alda score ≤3 (N=9)] are part of this analysis (clinical details in supplementary table 2). All DSM-IV psychiatric diagnoses were corroborated by two trained psychiatrists using the Mini International Neuropsychiatric Interview [9]. Healthy controls (N=12) with no axis I psychiatric illness and no family history of such illness in the past two generations were also recruited. The NIMHANS ethics committee approved the study protocols and informed written consent was obtained from all study participants.

**Figure 1:**
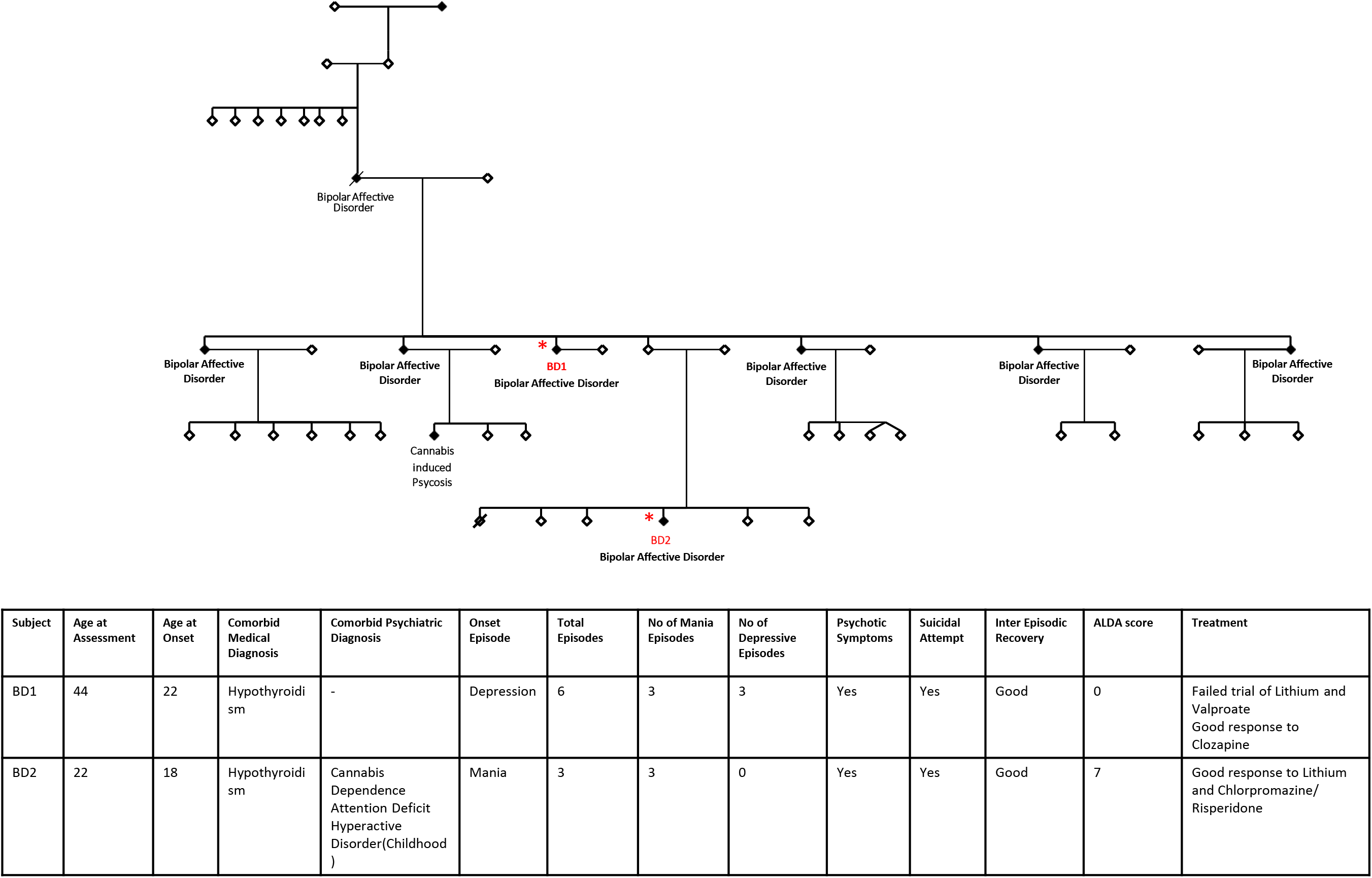
Family A pedigree with clinical details of BD1 (lithium non-responder) and BD2 (lithium responder).

### LCL generation and characterization

Lymphoblastoid cell lines were generated using Epstein Barr Virus from peripheral blood mononuclear cells as previously described [10].The cells were grown in RPMI-1640 (Himedia) medium containing 15% heat-inactivated fetal bovine serum (Gibco), 1% Penicillin-Streptomycin (Gibco) and 1% Glutamax (Gibco) as a suspension culture in 5% CO_2_ incubator at 37^0^C. Immunophenotyping of LCLs [11] by flow cytometry (BD FACSVerse, BD Biosciences, USA) confirmed that the cells were positive for B cell marker CD19 and negative for both, the T cell marker CD3 and the Natural Killer cell marker CD56 (supplementary figure 1A).

### Differentiation of NPCs from human IPSCs

Human PSCs were obtained from the ADBS [7] for two patients with BD (lines BD1 and BD2) in family A, and one unrelated healthy control (C1). The IPSCs had been generated from LCLs as previously described [12, 13]. Whole exome sequencing from this family has already been published [14] and rare damaging variants in BD1 and BD2 are shown in supplementary table 3. A fibroblast-derived control IPSC (C2) was also used for the experiments.

NPCs were generated as previously described [15]. A well-characterized high-quality IPSC culture was enzymatically dissociated using StemPro Accutase (Gibco) and cultured till day 7 in suspension in Embryoid body (EB) media [Knockout DMEM (Gibco), 20% KOSR (Gibco), 0.1mM Non-Essential Amino Acids (Gibco), 2mM Glutamax, 1% Penicillin-Streptomycin (Gibco), and 0.1mM Betamercaptoethanol (Gibco)]. EB media was replaced with Neural Induction Media [DMEM/F12 (Gibco), N2 supplement (Gibco), 8ng/ml bFGF (Gibco), 1x Glutamax (Gibco), 1x Penicillin-Streptomycin (Gibco), 1x Non-essential Amino Acids (Gibco) and 2µg/ml Heparin (Sigma)] from days 7 to 14. Subsequently, EBs were plated on Matrigel (Corning) coated dishes and allowed to form neural rosettes. Neural rosettes were passaged by manual selection and tertiary rosettes were dissociated mechanically by pipetting and plated as an NPC monolayer. Media was then replaced with Neural Expansion Medium [DMEM/F12 (Gibco), N2 supplement (Gibco), B27 supplement without Vitamin A (Gibco), 8ng/ml bFGF (Gibco), 1x Glutamax (Gibco), 1x Penicillin-Streptomycin (Gibco), 1x Non-essential Amino Acids (Gibco) and 2µg/ml Heparin (Sigma)]. Quantitative immunolabelling revealed comparable NPC differentiation from IPSC lines (Nestin: C1-99.2±0.4%, C2-99.3±0.4%, BD1-99.1±0.6%, BD2-99±0.5%; Pax6: C1-95.2±1.4%, C2, 94.2±2% BD1-94.8±2.2%, BD2-96.3±1.8%; Figure 2A).

**Figure 2:**
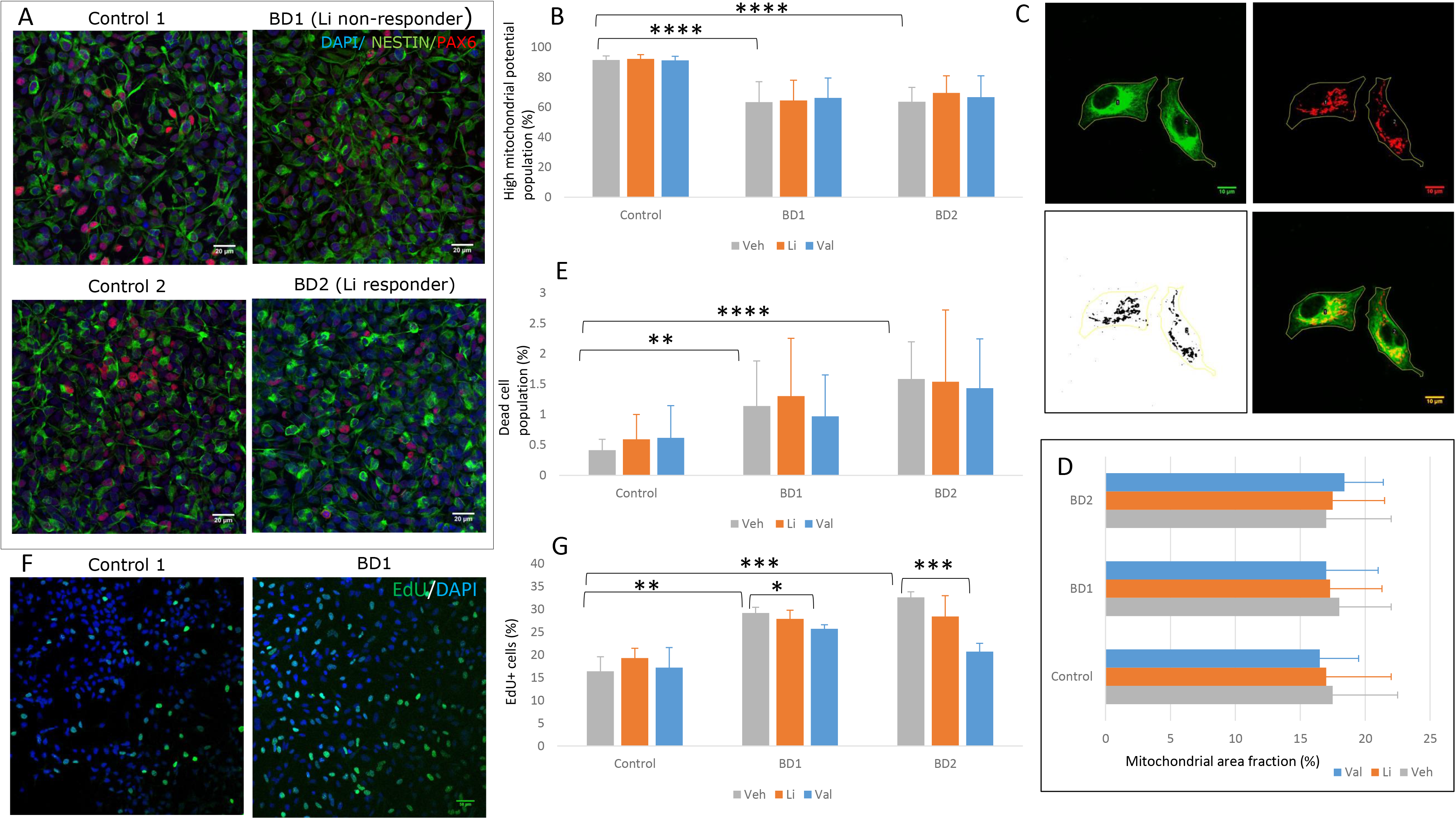
Experiments using NPCs. A) Representative immunocytochemistry of NPCs showing *Nestin+* cells in all the four cell lines used for experiments. Scale bar to be added. B) Comparison of high MMP population percentage (MTDR) in the three groups at baseline and after *in-vitro* treatment with lithium (1mM) or valproate (0.7mM) for 7 days by flow cytometry (N=5). C) Representative immunocytochemistry of NPCs showing *Nestin+* cells with mitochondria localized using *TOMM22*, and selection of regions of interest for calculation of mitochondrial area fraction. D) Comparison of mitochondrial area (N=3). E) Comparison of dead cell population percentage in the three groups at baseline and after *in-vitro* treatment by flow cytometry (N=5). F) Representative immunocytochemistry of NPCs showing *EdU+* cells indicating cells in S phase of cell cycle. G) Comparison of *EdU+* cells in the three groups at baseline and after *in-vitro* treatment (N=3). All data are shown as mean ± s.e.m.; Experiments were performed in at least 3 independent experiments (N) for each cell line; For group comparisons, initially appropriate ANOVA test was applied and if significant, independent multiple comparison test were performed. Significance level: *p≤0.05, **p ≤ 0.01, ***p ≤ 0.001, ****p ≤ 0.0001. Abbreviations: NPCs, neural precursor cells, MTDR, mitotracker deep red; MMP, mitochondrial membrane potential, EdU, 5-ethynyl-2’-deoxyuridine.

Gross chromosomal integrity checked by karyotyping was normal. Mycoplasma contamination was checked regularly with an enzyme-based mycoplasma detection kit (Lonza) as per manufacturer’s instructions.

### Drug treatment in NPCs

Cells were seeded at a density of 100 000 cells/cm^2^ on a tissue culture-treated surface additionally coated with Matrigel. Approximately 12 hours post-seeding; the drugs were added to the media as follows: Lithium Chloride (Sigma) at a working concentration of 1mM, Sodium Valproate (Sigma) at a working concentration of 0.7mM and phosphate buffer saline as the vehicle to the untreated control. Cellular assays and RNA extraction were performed on day 7.

### Drug treatment in LCLs

Four million LCLs were seeded in each of the three T25 flasks having RPMI-1640 complete media as described before with 1) vehicle (no treatment), 2) 1mM Lithium Chloride, and 3) 0.7mM Sodium Valproate. Post 7 days of treatment, LCLs were processed for cellular assays. Cells were also used for DNA and RNA extraction.

### Immunocytochemical analyses of NPCs

Cells were fixed in 4% paraformaldehyde (Sigma) for 20 min, permeabilized in 0.1% Triton X-100 (Invitrogen) at room temperature for 10 min and blocked in 3% donkey serum for 40 min. They were then incubated in primary antibodies for 60 min (Supplementary Table 6) followed by secondary antibodies for 30 min (Alexa Fluor dyes, Invitrogen; Supplementary Table 6). The nuclei were counterstained with DAPI (4’,6-diamidino-2-phenylindole, Life technologies) for 5min and coverslips were mounted on slides with Vectashield (Vector labs). Fluorescent imaging was performed on fields of view containing uniform DAPI staining using a Fluoview 3000 (Olympus) microscope. Images were processed with ImageJ64 (v 1.47) software and immunolabelled cells counted manually by a blinded observer. At least 10 representative images were taken for each experiment, which was done in triplicates.

### Whole transcriptome sequencing (RNA-Seq) and analysis

RNA-Seq was performed on the Illumina® Hi-Seq platform as per manufacturer’s protocol. There were two biological replicates available per sample. FastQC (v0.11.5; http://www.bioinformatics.babraham.ac.uk/projects/fastqc) was used for the quality of raw reads. It examines per base and per sequence quality scores, per base and per sequence GC content, per base N content and sequence length distribution. Cutadapt (v2.4) was used to remove adapter contamination in raw reads. The filtered reads were aligned to the human reference genome hg19 (GRCh37) using HISAT (v2.1). SAM to BAM conversion and sorting were done with Samtools 1.3 (https://sourceforge.net/projects/samtools/files/samtools/1.3/). The relative abundance of transcripts, measured as FPKM (Fragments Per Kilobase of transcript per Million mapped reads), was estimated using Cuffdiff (v2.02). Differentially expressed genes in response to lithium treatment were determined from the FPKM values obtained for each gene by calculating the fold change. Genes which showed >2-fold difference with FDR adjusted P-value <0.05 were considered as differentially expressed genes.

### Mitochondrial membrane potential (MMP) and cell death assay

Live staining with Mitotracker Deep Red (MTDR, Invitrogen, at a working concentration of 100nM) and a vital dye, Propidium Iodide (Invitrogen) at 15µg/ml in NPCs, Sytox Green (Invitrogen) at 30nM in LCLs followed by flow cytometry (BD FACSVerse™) was done to assess MMP and cell death respectively. Verapamil (Sigma) was also included at a working concentration of 5µM so as to prevent potential dye leak. The unstained tubes and single stained tubes were prepared similarly for the experiments to normalize before the analysis and avoid false positives. Experiments were performed in triplicates. FlowJo software was used for the analysis. Gates were applied on scatter plot using FSC vs SSC parameters to remove debris from the analysis. Finally, quadrant gates were applied using MTDR vs Sytox green signal to analyze the cell population of interest. The change in the percentage of the cell population of interest (gated in Q1 and Q3) was examined to study cell viability and MMP (supplementary figure 1B).

Pharmacological agents were included as a positive depolarization control. Treatment of cells with Carbonyl cyanide m-chlorophenyl hydrazone (CCCP) (Sigma), a respiratory uncoupler/ protonophore, at 50µM [16, 17] or 2% paraformaldehyde (Sigma) [18, 19] for 30 minutes resulted in significant reduction of MMP (supplementary figure 1C). This certifies the usage of MTDR to investigate MMP, as has been reported in previous studies [20–23]

### Mitochondrial area in NPCs

Immunocytochemistry and analysis were performed as described earlier. Individual regions of interest were first chosen based on Nestin staining and mitochondrial immunolabelling with anti-TOMM22 was assessed in binary images.

### Mitochondrial DNA content in LCLs

Total genomic DNA was extracted from LCLs and relative mitochondrial DNA (mtDNA) copy number was estimated by SYBR Green assay in a q-PCR system (Thermofisher). Cytochrome b (Cyt B) and NADH dehydrogenase 1 (ND1) genes were used to represent the mtDNA, and pyruvate kinase (PK) gene was used to represent the nuclear DNA. The primers selected for this experiment were based on an earlier study for mtDNA copy number by Gu et al., 2013 [24]. Relative mtDNA copy numbers of the genes Cyt B and ND1, normalized to the single-copy nuclear gene PK and relative to the calibrator is given by 2-ΔΔCt.

### Cell proliferation/ cycle assays

NPC proliferation assay was performed using the Click-it EdU Alexa Fluor 488 imaging kit (Invitrogen). EdU was added at a concentration of 10µM in media incubated for an hour at 37^0^C. Subsequently, cells were fixed and immunocytochemistry performed using the EdU detection cocktail and DAPI. The ratio of EDU positive to DAPI positive nuclei gave the percentage of cells in the proliferative phase.

Cell cycle assay was performed in LCLs fixed in 70% ethanol and subsequently incubated with Propidium Iodide dye (Invitrogen) at 15ug/ml and RNase A (Invitrogen) at 40ug/ml at 37^0^C for 30 min. Flow cytometry was performed as described earlier. FlowJo software was used for the analysis. Appropriate gates were applied on scatter plot using parameters required to remove debris and clumps from analysis while including only singlet cells. A histogram plot was generated for the gated singlet cells. The histogram plot was analyzed to estimate the percentage of cells in different phases of cell-cycle (i.e. G0/G1 or 2N phase, S phase and G2/M or 4N phase) (supplementary figures 1D, 1E).

### Gene expression studies in LCLs

To estimate the relative gene expression, total RNA was isolated using Trizol (Ambion) and converted to cDNA using SuperScript VILO cDNA Synthesis Kit (Thermofisher). The calibrator sample was used to calculate the relative quantification (RQ). The use of calibrator also normalizes inter plate assay variation for multiple runs.

The TaqMan gene expression assays were used for relative quantification of the target genes. *GAPDH* or *HPRT* were used as housekeeping genes. The q-PCR was duplexed by simultaneous amplification and quantification of both, target gene and HK gene in a single q-PCR reaction. The test sample cDNA along with No Template Control (NTC), calibrator, and Reverse Transcriptase (RT) negative sample were run in triplicates using the Real-time q-PCR system (AB7500; Thermofisher) to achieve the threshold cycle (Ct) value. The calculation used to estimate relative gene expression or RQ value is as follows:

ΔCt = Ct Target gene - Ct HK gene;

ΔΔCt = ΔCt Test sample - ΔCt Calibrator sample;

The relative gene expression or RQ value, normalized to an endogenous gene and relative to a calibrator, is given by 2 - ΔΔCt.

## Results

### RNA-Seq analysis of NPCs with and without in vitro lithium treatment

BD1 (Lithium non-responder), BD2 (Lithium responder) and control NPCs (C1 and C2) were treated *in vitro* with vehicle or lithium (1mM) for 7 days, followed by RNA-Seq analysis (supplementary table 4) to investigate molecular markers of drug response. Applying a false discovery rate of 0.05, there was no differentially expressed gene between NPCs, with and without *in vitro* lithium. Other comparisons were underpowered due to the genetic variability between the studied NPCs.

### Mitochondrial membrane potential (MMP) and cell viability assays in NPCs

We investigated whether there were deficits in MMP and cell viability in patient-derived NPCs. Reversal was attempted with lithium (1mM), and valproate (0.7mM). The doses used were based on the physiological dose range and available literature on cellular changes with *in vitro* drug treatment [24–26].

BD and control NPCs were treated *in vitro* with lithium/ valproate for 7 days, followed by flow cytometry analysis. At baseline, the percentage of cells with high MMP was found to be significantly lower in BD NPCs compared to control (Figure 2B). However, there were no significant differences between BD1 and BD2 or effect of *in vitro* drug treatment. Mitochondrial area fraction measured by TOMM22 immunolabeling was similar across all groups with or without drug addition (Figure 2C, 2D).

Baseline comparison for cell viability revealed, BD NPCs had a significantly higher percentage of dead cells compared to controls (Figure 2E). No significant differences were seen between BD1 and BD2, or with *in vitro* drug treatment.

### Cell proliferation assay in NPCs

Our experiments showed more EdU labelled cells (S phase) in BD NPCs compared to control (Figures 2F, 2G). As in the earlier assays, there were no significant differences between BD1 and BD2, or with in vitro drug treatment.

### Experimental validation of NPC results in larger sample size of LCLs

Our experiments showed clear cellular phenotypes in BD NPCs compared to controls; however, there was no reversal with *in vitro* drug treatment or effect of clinical lithium response status. As the objective is clinical translation, we decided to investigate these results in a peripheral model system. We used the LCL, which is an accepted cellular model for BD, from healthy controls and patients with extremes of lithium response.

As was evident in the NPCs, percentage of cells with high mitochondrial membrane potential was found to be significantly lower in BD LCLs compared to control (Figure 3A) with no significant differences between clinical lithium response groups. However, unlike the NPCs, *in vitro* lithium and valproate reversed the deficits in both groups of BD LCLs. Mitochondrial DNA copy number in LCLs measured by relative expressions of *ND1* and *CytB* was similar across groups (Figure 3B, 3C).

**Figure 3:**
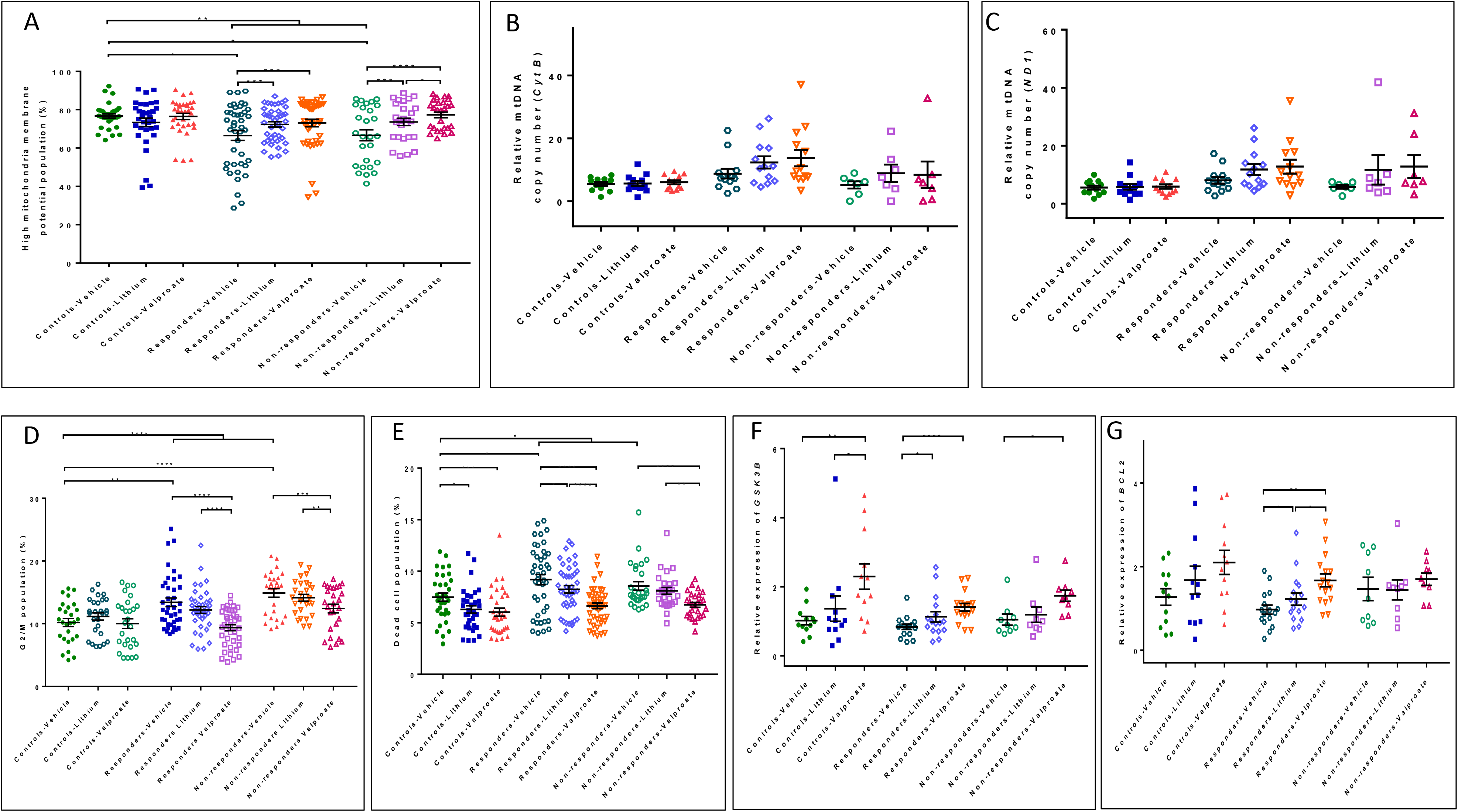
Experiments using LCLs. A) Comparison of high MMP population percentage in the three groups at baseline and after *in-vitro* treatment with lithium (1mM) or valproate (0.7mM) for 7 days. B) Comparison of relative quantification values of mitochondrial DNA-*Cyt B*, and C) *ND1*, from qPCR normalized to *PK (*single copy nuclear DNA) at baseline and after treatment with vehicle or lithium. D) Comparison of G2/M population percentage in the three groups at baseline and after *in-vitro* treatment. E) Comparison of dead cell population percentage in the three groups at baseline and after *in-vitro* treatment. F) Relative gene expression of *GSK3B* across groups - comparison for RQ expression values of *GSK3B* gene from qPCR normalized to *GAPDH* (endogenous control). G) Relative gene expression of *BCL2* across groups - comparison for RQ expression values of *BCL2* gene from qPCR normalized to *GAPDH* (endogenous control). All data are shown as mean ± s.e.m.; Experiments were performed in 3 independent experiments for each cell line; N, number of subject LCLs in each group. For group comparisons, initially appropriate ANOVA test was applied and if significant, independent multiple comparison test were performed. Significance level: *p≤0.05, **p ≤ 0.01,***p ≤ 0.001, ****p ≤ 0.0001. Abbreviations: LCLs, lymphoblastoid cell lines, MTDR, mitotracker deep red; MMP, mitochondrial membrane potential, PI, propidium iodide.

Similar to NPCs, BD LCLs had higher numbers of proliferative cells (G2/ M) in both clinical lithium response groups (Figure 3D). Valproate addition reversed the abnormality in both groups, but lithium addition did not.

In an interesting result, although the dead cell percentage was significantly higher in BD LCLs, reversal with *in vitro* lithium occurred only in the clinical lithium responder group (Figure 3E). Valproate reversed this deficit in both BD groups.

### Gene expression analysis related to cell viability

To identify specific molecular markers related to the cell viability phenotype, we performed LCL gene expression analysis of *BCL2*, *GSK3B* and *NR1D1*, all of which are involved in the mechanism of lithium action/ risk to bipolar disorder and cell survival (supplementary table 5). *BCL2* gene encodes an integral outer mitochondrial membrane protein that regulates cell death by controlling the mitochondrial membrane permeability [27]. *GSK3B* is involved in multiple signaling pathways controlling metabolism, differentiation, as well as death and survival [28]. *NR1D1*, also known as *Rev-ErbAα*, encodes a transcription factor that is a member of the nuclear receptor subfamily 1. Deletion of this gene affects the survival of neurons during postnatal development [29]. *In vitro* lithium specifically enhanced the expression of *BCL2* and *GSK3B* in LCLs from clinical lithium responders (Figure 3F, 3G), but not in other groups. *NR1D1* expression was unchanged across groups and on *in vitro* drug treatment.

## Discussion

In this study, we aimed to identify cellular models for BD and response to lithium treatment. Not all BD patients respond favorably to lithium, though percentages vary across studies. This makes it pertinent to identify molecular markers associated with clinical response to lithium, given that it is the primary drug of choice for BD.

Both the occurrence of disease (BD) and response to treatment are influenced by genetic factors. Therefore, to delineate the mechanisms that could have a bearing on response to lithium, we chose two BD patients from the same family (i.e. shared genetic background) who were discordant in their response to lithium (Fig. 1) and characterized the cellular phenotypes for disease and lithium treatment.

Mitochondrial dysfunction is one of the most consistently replicated findings in the literature on BD [30, 31]. Consistent with published data, we also find that the percentage of cells with high MMP is reduced in BD patients in our cohort (Figs. 2B, 3A). Independent exposure *in vitro* to either lithium or valproate increased this value in LCLs (Fig. 3A) but not in patient-derived NPCs (Fig. 2B). Previous studies have shown that lithium treatment increased mitochondrial complex activities of leukocytes [32] and that valproate treatment enhanced *NDUFV2* gene expression [33].

The percentage of proliferating NPCs is higher in BD patient derived cells (Fig. 2G) as well as LCLs (Fig. 3D). Presence of the same phenotype in NPCs and LCLs in BD patients strongly argues in favor of increased cell proliferation as a marker associated with the disease. Our conclusion is supported by previous studies [34–37] that have reported dysregulation of genes associated with G1 or G2 checkpoint of the cell cycle in samples from BD patients. We also found that exposure to lithium *in vitro* did not make a difference in this phenotype, neither in LCLs nor in NPCs; even when patients did show favorable response to clinical lithium treatment. Similar discordance between clinical response to lithium and *in vitro* exposure of patient-derived NPCs has also previously been reported by Squassina and colleagues (2016) [38], and by Pietruczuk et al., (2018) [39] on patient-derived T-lymphocytes. Collectively, these data demonstrate that cell proliferation is a phenotype that strongly correlates with disease, but is not a reliable marker of clinical response to lithium.

In contrast to the response seen in NPCs, we found that, in LCLs from a larger group of BD patients, *in vitro* lithium treatment helps maintain viability as demonstrated by a decrease in the percentage of dead cells (“Responders-Lithium,” Fig. 3E). From its clinical use in BD over sixty years ago [40], several molecular pathways have been implicated in the benefit conferred by lithium [41]. Included in this shortlist of cellular parameters is cell viability [42, 43]. The question then is to identify a convenient cellular model which is indicative of clinical efficacy of lithium. Data presented here argue in support of LCLs being such a model system.

GSK3B, a direct target of lithium, modulates various functions including growth metabolism, cell signaling and multitude transcription factors that in turn regulate cell survival [28,44,45]. Comparable to our study design, Geoffrey et al. showed *in vitro* lithium (at 1mM) exposure for eight days increased *GSK3B* mRNA [46]. Though we observed similar trends on *GSK3B* expression levels, a statistically significant increase was evident only in clinical lithium responders. Shorter durations of lithium exposure have not shown *GSK3B* expression differences in earlier studies [46, 47].

We also report here an increase in *BCL2* expression concomitant with *in vitro* exposure to lithium in LCLs (Fig. 3G). This is significant because lymphocytes derived from BD patients are more prone to apoptosis, with increased expression of the pro-apoptotic BAX protein [39]. In this context, increased levels of anti-apoptotic *BCL2*, evident exclusively in cells derived from BD patients who clinically responded to lithium; strongly relates cell viability as a proxy marker for clinical response to lithium.

We have recently reported on clinical factors associated with lithium response in BD [5]. In the present study, we intended to extend our previously reported findings, in an attempt to relate clinical factors with cellular parameters associated with BD and its treatment with lithium. We report here that the clinical effectiveness of lithium correlates well with increased cell viability in LCLs. Currently, we are investigating whether the same cellular parameter i.e. cell viability can also predict clinical responsiveness to lithium in lymphocytes from treatment-naive BD patients, who will be in prospective clinical follow-up after initiation of lithium.

## Acknowledgements

The authors would like to thank Dr. Preeti Joshi, (Retd.) Professor of Biophysics, NIMHANS; Dr. Y C Janardhan Reddy, Professor of Psychiatry, NIMHANS and Dr. Mitradas M. Panicker (Retd.) Faculty of Neurobiology, NCBS, for providing critical inputs during various parts of the project. We are grateful towards Dr Manjunath, Assistant Professor of Neurovirology, NIMHANS and Phillippo, PhD Scholar, NIMHANS for their constructive criticism and valuable suggestions related to FACS experiment and also allowing us to access the Flowcytometry facility. We would also like to thank Ms. Varalakshmi R. and Mr. Suneela Kumar B. for the technical support. We would like to thank the clinicians and staff at the NIMHANS; as well as the subjects and their families for their co-operation.

