## supplementary figure 1 for "Studying cellular functions in bipolar disorder: Are there specific predictors of lithium response?"

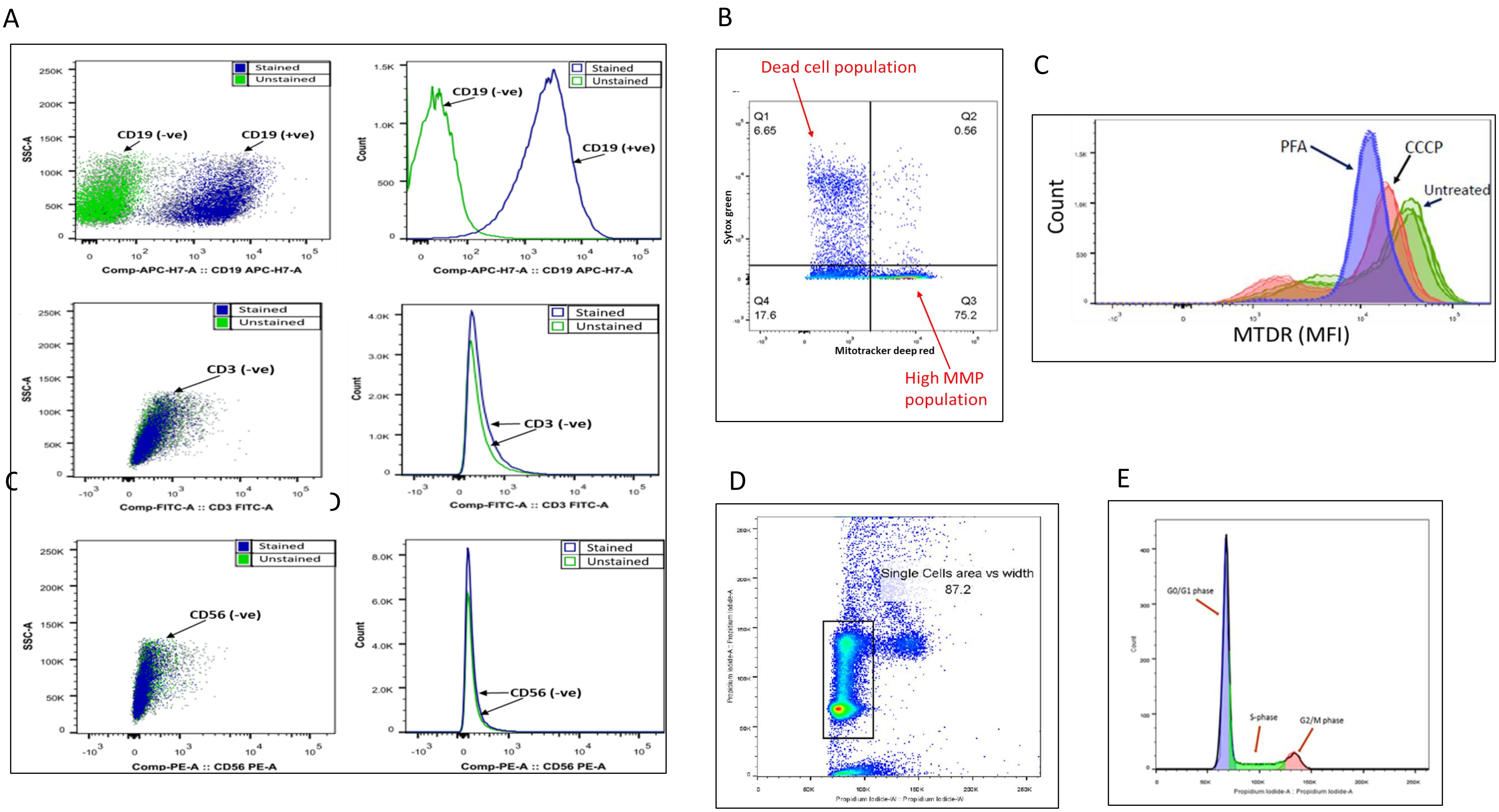
A
Stained
Unstained
CD19 (-ve)
CD19 (+ve)
SSC-A
Count
Comp-APC-H7-A :: CD19 APC-H7-A
CD3 (-ve)
Comp-FITC-A :: CD3 FITC-A
CD56 (-ve)
Comp-PE-A :: CD56 PE-A
B
Dead cell population
Q1
6.65
Q2
0.56
Q4
17.6
Q3
75.2
Sytox green
Mitotracker deep red
High MMP population
C
PFA
CCCP
Untreated
Count
MTDR (MFI)
D
Single Cells area vs width
87.2
E
G0/G1 phase
S-phase
G2/M phase

Supplementary figure 1. A) Characterization of LCLs by immunophenotyping- Scatterplot and histogram plot showing LCLs positive for *CD19* (B cell marker) (top panel), negative for *CD3* (T cell marker) (middle panel), negative for *CD56* (NK cells) (bottom panel). B) Representative flow cytometer scatter plot for mitochondrial potential and cell viability assay**.** Dot plot for MTDR signal against Sytox Green, further gated into four quadrants based on live or dead cells (Sytox positive indicate dead cells) and high or low MMP population (based on MTDR signal intensity). The percentage of cells in Q1 (dead cell) and Q3 (live cells with high MMP) were analyzed further to assess cell viability and MMP respectively. C) Mitochondrial membrane depolarization positive control experiment- Histogram plot showing the change in mean MFI of MTDR after incubation with CCCP (50uM) and PFA (2%). D) Representative flow cytometer scatter plot for cell cycle assay. Dot plot for PI**,** to show gating of single cells using width versus area parameters. E) Illustrates PI area parameter histogram plot of the singlet cells to determine percentage of cells in G0/G1, S and G2/M phases of the cell cycle using FlowJo software. Abbreviations: LCLs, lymphoblastoid cell lines, PBMCs, peripheral blood mononuclear cells, EBV, Epstein-barr virus, MFI, mean fluorescence intensity, MTDR, mitotracker deep red, CCCP, carbonyl cyanide m-chlorophenyl hydrazine, PFA, paraformaldehyde.
