## supplementary table 3 for "Studying cellular functions in bipolar disorder: Are there specific predictors of lithium response?"

**Supplementary table 3: Rare damaging exome variants identified in family A**

| Gene | Variant | RS id/Novel | No of affected | Presence in BD1/BD2 | Signaling pathway | Cellular role |
|---|---|---|---|---|---|---|
| **DENND5A** | c.A2699G | rs779817963 | 4 | BD1/BD2 | ERK pathway [1] | Cell migration [2], proliferation, apoptosis |
| **KIF7** | c.G2690C | rs749711306 | 3 | BD1 | Hedgehog signalling [3,4] | Cell proliferation [5], migration |
| **SCN3A** | c.G83A | rs775711350 | 3 | BD1/BD2 | No pathway reported | Neuronal migration [6], Cell cycle [7] |
| **PARP14** | c.G3467A | Novel | 4 | BD1/BD2 | JNK2 signalling [8] | Apoptosis [9], glycolysis [10] |
| **PCCB** | c.C595T | rs371155999 | 3 | BD1/BD2 | Proprionate metabolism pathway [11] | Mitochondrial oxidative phosphorylation [12] |
| **TRMT44** | c.C1405T | rs373816157 | 3 | BD1 | | |
| **NRG2** | c.C1477T | rs148371256 | 3 | BD1 | MAPK and ERBB pathway [13,14] | Cell migration, proliferation [15,16], oxidative stress [17] |
| **NIPBL** | c.A4496C | Novel | 4 | BD1/BD2 | Notch pathway [18], Cohesion [19], Wnt and PI3K-AKT pathway [18] | Cell migration, proliferation, apoptosis [20,21] |
| **SCUBE3** | c.C1996T | Novel | 3 | BD1/BD2 | FGF Pathway [22] and hedgehog signaling [23] | Cell proliferation [24] |
| **ANLN** | c.C128T | rs575071809 | 3 | BD1/BD2 | PI3K pathway [25] | Cell migration, Cell cycle [26,27] |

12. Chapman KA, Ostrovsky J, Rao M, Dingley SD, Polyak E, Yudkoff M, et al. Propionyl-CoA carboxylase pcca-1 and pccb-1

gene deletions in Caenorhabditis elegans globally impair mitochondrial energy metabolism. J Inherit Metab Dis. 2018;41:157–168.
