## supplementary table 5 for "Studying cellular functions in bipolar disorder: Are there specific predictors of lithium response?"

**Supplementary table 5: Studies of *BCL2. GSK3B and NR1D1* genes in bipolar disorder**

| Author and Year | Type of study | Sample details | Objective of study | Significant results |
|---|---|---|---|---|
| | | ***BCL2*** | | |
| H.-W. Kim, Rapoport, & Rao, 2010 [1] | Case- control study (protein/ mRNA level) | 10BD &10 control frontal cortex from PM brains | To investigate levels of BCL2 protein & mRNA in BD compared to HC | Decreased BCL2 protein and mRNA levels in frontal cortex from BD brain tissues compared to controls.<br>BAX/BCL2 ratio was increased in BD brain tissues. |
| Moutsatsou et al., 2014 [2] | Case- control study (mRNA level & apoptotic activity) | Lymphocyte from 35 BD and 10 HC | To investigate the level of *BAX/BCL2* mRNA ratio level, caspae3 activity and cytochrome C release in BD and controls. | Higher *BAX/BCL2* mRNA levels in BD patients in manic and depressed state compared to HCs. Cytochrome c release, caspase-3 activity was increased in manic and depressed BD patients compared to HC, indicating higher apoptotic activity in BD. |
| W. T. Chen, Huang, & Tsai, 2015 [3] | Case- control study (protein level) | 20 BD patients in manic phase and 40 HC from Taiwan | To examine the serum BCL2 levels in BD and HCs | Serum BCL2 levels higher in manic state of BD patients than HC, though statistically not significant. |
| Uemura et al., 2011 [4] | Functional study and case- control association study | LCLs from 245 patients (150 BD-I, 65 BD-II &30 MDD) and 70 HC subjects | To study the role of *BCL2* rs956572 SNP on basal intra cellular calcium, mRNA and protein levels in BD and control subjects. . | 1) No significant association of the *BCL2* SNP with any of the disorders.<br>2) Basal calcium levels of LCLs: Significantly higher in BD compared to controls, subjects with GG genotype being the highest compared to other genotype (AA<AG<GG). BD with GG reported to have higher levels compared to other groups carrying same GG.<br>3) *BCL2* mRNA and protein levels were lower in BD than HC. GG genotype subjects lower levels compared to AA or AG, effect was more prominent in BD. |

| | | | | |
|---|---|---|---|---|
| Soeiro-de-Souza et al., 2013 [5] | Case-control association study | 40 BD euthymic patients and 40 HC from Brazil. MRS is done to obtain glutamate levels of anterior cingulate cortex (ACC) | Tested the association of *BCL2* (rs956572) SNP with ACC glutamate levels | AA genotype at the *BCL2* SNP was associated with elevated ACC glutamate metabolites in the BD patients, not in controls. |
| Uemura, Green, & Warsh, 2015 [6] | Functional study and case- control association study | LCLs derived from 215 BD cases, including 150 BD-I and 65 BD-II; and 70 healthy controls | To investigate whether the *BCL2* rs956572 variant associates with intracellular calcium dyshomeostasis in BD and HC | 1) Lower $Ca2^{+}$ in subjects with BCL2 rs956572 AA variant compared to AG or GG as a whole.<br>2) BD patients carrying GG genotype- highest $Ca2^{+}$ compared to AA or AG.<br>3) BD patients carrying GG genotype- higher $Ca^{2+}$ compared to HC with GG genotype. |
| Corson, Woo, Li, & Warsh, 2004 [7] | Pharmacological study | Human hNT neurons from NT2 teratoma cells and SVG p12 SV40 glia cells (treated with lithium or valproate for 7days) | To test the effect of lithium and valproate on *BCL2* mRNA levels in hNT neurons | Treatment of hNT cells with valproate (0.35, 0.75, 1 mM) for 7 days upregulated BCL2 mRNA (max increase with 0.75mM), whereas lithium (0.75-2mM) could not alter. The SVG glia cells *BCL2* mRNA was not changed by both the treatments. |
| Lai et al., 2006 [8] | Pharmacological study | Human SH-SY5Y neuroblastoma, SVGp12 glial cells and U87 glioma cells (treated with lithium [1mM] or valproate [0.6mM] for 1 or 7 days) | Looked into the effect of lithium or valproate in stress induced human cells | Pretreatment of SH-SY5Y cells for 7 days with lithium or valproate significantly reduced rotenone or $H_2O_2$ induced cytochrome C release, caspase activity & cytotoxicity and upregulated BCL2 protein level. No effect was reported on 1 day treatment with the lithium or valproate. Other cell types were not affected by any such treatments. |
| Creson, Yuan, Manji, & Chen, 2009 [9] | Pharmacological study | Human SH-SY5Y neuroblastoma cells | Looked in the effect of valproate on BCL2 protein and mRNA levels in human neuroblastoma cells. | Increased BCL2 protein in concentration (0.125-2mM) & time (1-3 days on 0.8mM) dependent manner. 0.5-2mM for concentration and 2 or 3 days treatment induced the increase in BCL2 level whereas 1 day did not. Similarly, valproate increased the BCL2 mRNA in time dependent manner, 3rd day being the highest increase. The enhancement of BCL2 levels was selective as the HK gene GAPDH was not altered. |

| | | | | |
|---|---|---|---|---|
| Odeya D, Galila A, & Lilah T, 2018 [10] | Pharmacological study | 10-12 weeks old wild type mice from *IMPA1* colony | Effect of lithium on *BCL2* gene expression in hippocampi of mice | On treatment with lithium *BCL2* gene expression levels were increase when normalized to *ACTB*, whereas decreased on normalizing with *MAPK6* or *ANKRD11*. |
| Machado-Vieira et al., 2011 [11] | Functional study- 1) SNP- with intracellular calcium, mRNA & protein, 2)SNP-lithium effect | LCLs from 18 BD subjects with equal numbers of individuals carrying the rs956572 variants for AA, AG & GG | To study the role of *BCL2* rs956572 SNP on basal intra cellular calcium and after 1mmol/L lithium treatment in BD LCLs for 7 days. | 1) Basal and stimulated intracellular $Ca2^+$ were higher in BD LCLs with AA variant compared to GG variant though, BCL2 mRNA and protein level was least in AA variant of the *BCL2* polymorphism.<br>2) Li treatment increased the *BCL2* expression in the AA variants of BD LCLs. |
| Lowthert et al., 2012 [12] | Pharmacogenetic study | 8 weeks of open label lithium study in 20 BD patients | To assess the change in gene expression in peripheral blood of lithium responders and non-responders | 127 genes differentially expressed between responders and non-responders.<br>Pathway analysis showed regulation of apoptosis was significantly affected.<br>Upregulation of anti-apoptotic gene *BCL2* & downregulation of pro-apoptotic genes (*BAD*, *BAK1*) in responders and inverse relation in NR after 4 weeks. |
| ***GSK3B*** | | | | |
| Xiaohong Li, Liu, Cai, Wang, & Li, 2010 [13] | Case- control study and anti-manic treatment response (protein level) | 1) 30 medication free BD manic subjects were compared with 30 healthy controls from Beijing Anding Hospital, China.<br>2) 47 BD (4 weeks) and 28 BD (8 weeks) subjects were analyzed for pre and post treatment with lithium, valproate and atypical anti-psychotics | To test the regulation of GSK3B in BD patients with manic episode and in response to treatment - examined the protein level and the inhibitory serine phosphorylation of GSK3B in PBMCs of patients compared with healthy controls | 1) The total protein levels of GSK3B was significantly higher in BD manic subjects than in healthy controls.<br>2) Phospho-Ser9-GSK3B was reported to be trend toward lower in bipolar manic subjects than in healthy controls.<br>3) Significant increase in phospho-Ser9-GSK3B was reported in 28 BD subjects post 4 weeks and 8 weeks treatment.<br>4) Total GSK3B among the 47 subjects was not significantly different at treatment. |

| | | | | |
|---|---|---|---|---|
| Pandey, Ren, Rizavi, & Dwivedi, 2010 [14] | Case- control study (protein level) | 1) 21 BD patients and 21 HC from Chicago were investigated for GSK3B level in platelet,<br>2) The patients compared for GSK3B level before and after treatment (lithium or valproate or antipsychotics) for 8 weeks. | To explore the role of GSK3B in BD and in response to treatment | 1) GSK3B protein level in cytosol and membrane fraction of platelets from BD were decreased in comparison to controls.<br>2) The protein level after 8 weeks of treatment was increased. |
| Lesort, Greendorfer, Johnson, 1999 [15] | Case- control study in PM brain tissues | DLPFC of 5 BD and 5 controls from Ohio. | To compare the levels of GSK3B protein in brain tissues of BD and HCs | No significant difference was reported. |
| Munkholm, Peijs, Vinberg, & Kessing, 2015 [16] | Case-control candidate gene expression study | 37 rapid cycling BD patients and 40 HC of Danish population | Tested *GSK3B* gene expression in BD & HC | *GSK3B* mRNA level was significantly downregulated in BD, however after Bonferroni correction the result was not significant. |
| Benedetti, Bernasconi, et al., 2004 [17] | Candidate gene association study | 185 Italian BD patients | To test the effect of the *GSK3B* rs334558 SNP on age at onset of BD | Homozygote TT was reported to be associated with earlier age at onset of BD. |
| Benedetti, Serretti, et al., 2004 [18] | Candidate gene association study | 60 depressed BD patients | To test the effect of *GSK3B* -50T/C polymorphism on age at onset of BD and acute response to total sleep deprivation | Homozygotes for the mutant allele was associated with later age at onset of BD, less severe symptomatology when depressed (HDRS score), & better acute effects of total sleep deprivation treatment on perceived mood (VAS score). |
| Nishiguchi, Breen, Russ, St Clair, & Collier, 2006 [19] | Case-control candidate gene association study | 280 Caucasian BD patients and 407 HC. | To test the association of the *GSK3B* -50T/C SNP and BD. | No significant association |

| | | | | |
|---|---|---|---|---|
| Szczepankiewicz, Skibinska, et al., 2006 [20] | Case-control candidate gene association study | 416 Polish patients and 408 HC | To test the association of the *GSK3B* -50T/C SNP and BD. | 1) Trend association of heterozygous T/C genotype with BD was reported.<br>2) Significant association of SNP with the female BDII patients (n=57).<br>3) No significant association with AAO of BD |
| Serretti et al., 2008 [21] | Candidate gene association study | 365 Italian mood disorder patients, included 122 MDD and 243 BD patients. | To evaluate the association of the polymorphism with symptomatic and personality feature in mood disorder | The GSK3B polymorphism was found associated with delusional symptomatology and with the personality features linked to self-transcendence. |
| Subhashree et al., 2009 [22] | Case-control association study (NIMHANS) | 186 subjects with BD and 186 healthy controls from NIMHANS, Bangalore, INDIA | To investigate the association of -50T/C SNP in *GSK3B* gene with BD. | No significant association |
| E. Jiménez et al., 2013 [23] | Candidate gene association study | 192 Caucasian BD subjects included 66 suicide attempters and 126 suicide non attempters | To investigate association between the *GSK3B* -50T/C SNP & suicide behavior in BD | C allele of the *GSK3B* SNP showed trend association (p=0.052) with suicide attempters. |
| Esther Jiménez et al., 2014 [24] | Candidate gene association study | 199 Caucasian BD subjects | To evaluate the effect of the SNP on impulsivity in BD | C allele carrier was associated with higher level of impulsivity in BD |
| Tang et al., 2013 [25] | Meta-analysis study | 48 relevant studies screened and 5 BD studies (3 Asian and 2 Caucasian) finally included for analysis. Total 971 cases and 1397 controls | To test the association of the *GSK3B* -50T/C SNP and BD. | No significant association of the *GSK3B* -50T/C SNP and BD. |
| G. Chen et al., 2014 [26] | Meta-analysis study | 95 relevant studies screened and 10 BD studies included<br>1) For association with BD | To investigate the association between the *GSK3B* -50T/C SNP and the susceptibility or age at | No significant association of the GSK3B -50T/C SNP with risk of BD or age at onset of BD was reported in any of the genetic model that was analyzed. |

| | | | | |
|---|---|---|---|---|
| | | risk, 6 (3 Asian and 3 Caucasian) publications. Total of 1251 cases & 1804 controls.<br>2) For association with age at onset, 6 (2 Asian and 4 Caucasian) publications were analyzed. Total of 659 cases. | onset of BD. | |
| De Sarno, Li, & Jope, 2002 [27] | Pharmacological study | 1) *In vitro*: Human neuroblastoma SH-SY5Y cells- with valproate or lithium for 24 hours at varying concentration<br>2) *In vivo*: Adult male C57BL/6 mice treated with 0.4% lithium (0.7mM at serum) | Evaluated the effect of lithium or valproate on phosphorylation of GSK3B. | 1) Sodium valproate treatment effected gradual increase in the inhibition-associated phospho-Ser9-GSK3B.<br>2) Lithium treatment increased the phospho-Ser9-GSK3B both in cells and in mouse brain after chronic administration. |
| Zhang, Phiel, Spece, Gurvich, & Klein, 2003 [28] | Pharmacological study | 293T cells, Neuro2A, and NIH3T3 cells from American Type Culture Collection- treated with lithium or VPA (varying concentration and duration) | Evaluated the effect of lithium or valproate on GSK3B. | 1) Lithium treatment inhibited GSK3B activity which was mediated through increased phospho-Ser9-GSK3Bin cells.<br>2) VPA was not reported to inhibit GSK3B & did not induce increase in phosphorylation of GSK3B. |
| Jonathan Ryves, Dalton, Harwood, & Williams, 2005 [29] | Pharmacological study- using rat primary cells | Rat neocortical neurons – treated with 3mM lithium or 1.8mM valproic acid (VPA). | To examine the effect of lithium and VPA on GSK3B protein levels. | 1) No effect of lithium and VPA on GSK3B protein level. |
| Abdul A, De Silva B, & Gary R, 2018 [30] | Pharmacological study | NIH-3T3 mouse embryo fibroblast, A172 human glioblastoma | Evaluated the effect of lithium or beryllium on phosphorylation of GSK3B and its substrate. | Lithium (20mM) and beryllium (30 & 100 uM) decreases phosphorylation of glycogen synthase (GS) and increased phosphorylation of Ser9-GSK3B in NIH3T3. No change in levels of total GS or GSK3B. |

| | | | | |
|---|---|---|---|---|
| | | | | In A172 cells, lithium increased phosphor-Ser33/Ser37-B catenin. |
| Xiaohua Li et al., 2007 [31] | Case-control study (protein level) using PBMCs of subjects and *in vitro* study by lithium treatment of PBMCs | 23 HC, 9 lithium treated BD and 13 lithium free BD subjects of Caucasian population. | Evaluated change in serine phosphorylation of GSK3B in PBMCs of BD and HC subjects at baseline and *in vitro* treatment with lithium (20mMol/L) for 1 hour | 1) Basal level of phospho-Ser9-GSK3B was reported to be lowest in HC followed by 3-fold increase in lithium free BD and highest in lithium treated BD subjects.<br>2) *In vitro* lithium treatment was also associated with elevation of phospho-Ser9-GSK3B level.<br>3) No change in total GSK3B protein level. |
| Mendes et al., 2009 [32] | Animal model study (gene expression)– lithium treatment using Wistar rat | 1) *In vitro*: cortical and hippocampal neurons- 5days LiCl treatment (0.02 to 2mM)<br>2) *In vivo*: 12 rats- (0.12mmol lithium; 0.24mmol lithium) | To test the role of role of *GSK3B* in response to lithium treatment. | 1) *In vitro*: *GSK3B* mRNA level was reduced in hippocampal neurons on treatment but no changes in cortical neurons.<br>2) *In vivo*: *GSK3B* mRNA reduced in hippocampus but not in cortex or in leukocyte of treated rats. |
| McCarthy et al., 2011 [33] | Pharmacogenetic study- gene expression in LCLs | LCLs from BD lithium responders (N=13) and lithium non-responders (N=18) | To test the effect of lithium 1mM for 72 hours on *GSK3B* expression in LCLS from BD lithium response patients | No effect on *GSK3B* gene expression in both the BD lithium response groups |
| Geoffroy et al., 2017 [34] | Pharmacogenetic study- gene expression in LCLs | 38 French Caucasian BD patients which included 16 ER and 20 NR of lithium. | To test the effect of lithium 1mM (2 -8 days) on *GSK3B* expression in LCLS from BD lithium response patients | Only on day 8 lithium significantly increased *GSK3B* gene expression in BD lithium non-responders |
| Benedetti et al., 2005 [35] | Pharmacogenetic association study | 88 BD patients- 2 years on lithium. | To test the association of the *GSK3B* -50T/C SNP with therapeutic response to lithium. | Mutant allele C carriers improved recurrent rate of mood episode after 2 years on lithium treatment. |
| Szczepankiewicz, | Pharmacogenetic association study | 89 polish BD patients-5 years on lithium | To test the association of G*SK3B* -50T/C SNP with | No significant differences in genotypic and allelic frequencies between the SNP |

| | | | | |
|---|---|---|---|---|
| Rybakowski, et al., 2006 [36] | | | therapeutic response to lithium. | and the degree of lithium response . |
| Numajiri et al., 2012 [37] | Pharmacogenetic association study | 29 Japanese BD patients | To test the association of the SNP with therapeutic response to lithium. | T allele significantly associated with lithium responders. |
| Y. F. Lin, Huang, & Liu, 2013 [38] | Case-control and Pharmacogenetic association study | 138 Taiwanese BD patients and 131 controls. 83 patients out of 138 cases were evaluated for lithium treatment (24 months) efficacy. | To test the association of the SNP with BD risk and therapeutic response to lithium treatment. | 1) No significant association of the *GSK3B* -50T/C SNP and BD.<br>2) TT genotype was associated with poor lithium treatment response. |
| Iwahashi et al., 2014 [39] | Pharmacogenetic association study | 42 Japanese patients: 27 were lithium responders and 15 were non-responders. | To test the association of SNP with lithium treatment response. | No significant difference was reported in genotype and allele frequency of the SNP between lithium responders and non-responders.<br>However, haplotype blocks T-A and C-A (with another SNP [-1727A/T]) was reported to be associated with higher lithium response and lower lithium response respectively. |
| Mitjans et al., 2015 [40] | Pharmacogenetic association study | Total 131 BD patients from Barcelona which included 26 excellent responders (ER); 62 partial responders (PR) and 43 non-responders (NR) based on lithium response. | To test the association of SNP with lithium treatment response. | No significant difference was reported in genotype and allele distribution between the lithium response groups.<br>However haplotype rs1732170-rs11921360-rs34558 was associated with lithium response. The C-C-A haploblock was significantly less frequent in group of lithium partial and non-responders than excellent responders. |
| | | ***NR1D1*** | | |
| Yang, Van Dongen, Wang, | Case-control association study | 2 set of fibroblast samples from Corriel Cell | To study the expression of core clock genes in BD and | Set-I: No difference in circadian period.<br>Amplitude of rhythmic expression of *NR1D1* |

| | | | | |
|---|---|---|---|---|
| Berrettini, & Bućan, 2009 [41] | | Repositories. Set-I: 12BD &12 HC; Set-II: 18BD & 35HC. | HC. | reduced in BD. Overall expression of *NR1D1* reduced in BD though statistically not significant.<br>Set-II: *GSK3B* mRNA & protein level no significant difference between BD and HC, whereas serine-9-phospho GSK3B was significantly reduced in BD. |
| Nováková, Praško, Látalová, Sládek, & Sumová, 2015 [42] | Case-control association study | Buccal cells from 19 HC, 22 BD depressive & 19 BD in manic subjects from Czech Republic. | To investigate the *NR1D1* expression profiling for 24 hours in buccal cells of BD and HC | *NR1D1* expression profiling for 24 hours in buccal cells. *NR1D1* expression profiles of BD in mania was advanced compared to depression and trend advanced compared to control. Amplitude *NR1D1* expression higher in mania. |
| Warburton et al., 2015 [43] | Pharmacological study | Human SH-SY5Y neuroblastoma cells | To examine the expression profiling in neuroblastoma cells on treatment with lithium (1mM) or valproate (5mM) for 1 hr. | Lithium treatment of neuroblastoma cells showed trend change in *NR1D1* expression, whereas valproate did not alter the expression. |
| McCarthy et al., 2011 [33] | Pharmacogenetic study | Genetic association: 282 BD Caucasian origin (148 lithium responders and 134 non-responders.<br>LCL experiment: 13 responders and 18 non-responders (1mM lithium for 72hours treatment) | To test the role of NR1D1 lithium treatment response. | 1) Allele A at rs2071427 SNP associated with lithium good response.<br>2) Homozygous allele A at the SNP decreased *NR1D1* mRNA (full length transcript) after lithium treatment compared to homozygous G allele.<br>3) AA genotype was associated with trend increase of *NR1D1* (both full and truncated transcript) mRNA compared to GG after treatment. |
| Geoffroy et al., 2017 [34] | Pharmacogenetic study | LCLs from 36 BD subjects (20 lithium responders and 16 non-responders of Caucasian origin | To analyse the gene expression of BD LCLs at day2,4,8 on 1mM lithium treatment | *NR1D1* gene expression was downregulated at day 2 in responders. *NR1D1* was upregulated at day 4 for both responders and non-responders. No significant changes at day 8. |

| | | | | |
|---|---|---|---|---|
| Campos-de-Sousa et al., 2010 [44]; Kishi et al., 2008 [45]; Kripke, Nievergelt, Joo, Shekhtman, & Kelsoe, 2009 [46]; Severino et al., 2009 [47]. | Case-control candidate gene association studies | BD and HC subjects in various studies | Tested the association of different *NR1D1* polymorphisms with risk of BD | The studies have reported positive association of the *NR1D1* polymorphisms with risk for BD. |

21. Serretti A, Benedetti F, Mandelli L, Calati R, Caneva B, Lorenzi C, et al. Association between GSK-3β -50T/C polymorphism and personality

and psychotic symptoms in mood disorders. Psychiatry Res. 2008;158:132–140.

30. Abdul AURM, De Silva B, Gary RK. The GSK3 kinase inhibitor lithium produces unexpected hyperphosphorylation of β-catenin, a GSK3 substrate, in human glioblastoma cells. Biol Open. 2018;7.

31. Li X, Friedman AB, Zhu W, Wang L, Boswell S, May RS, et al. Lithium Regulates Glycogen Synthase Kinase-3β in Human Peripheral Blood Mononuclear Cells: Implication in the Treatment of Bipolar Disorder. Biol Psychiatry. 2007;61:216–222.

32. Mendes CT, Mury FB, de Sá Moreira E, Alberto FL, Forlenza OV, Dias-Neto E, et al. Lithium reduces Gsk3b mRNA levels: implications for Alzheimer Disease. Eur Arch Psychiatry Clin Neurosci. 2009;259:16–22.

33. McCarthy MJ, Nievergelt CM, Shekhtman T, Kripke DF, Welsh DK, Kelsoe JR. Functional genetic variation in the Rev-Erbα pathway and lithium response in the treatment of bipolar disorder. Genes Brain Behav. 2011;10:852–861.

34. Geoffroy PA, Curis E, Courtin C, Moreira J, Morvillers T, Etain B, et al. Lithium response in bipolar disorders and core clock genes

expression. World J Biol Psychiatry. 2017:1–14.

47. Severino G, Manchia M, Contu P, Squassina A, Lampus S, Ardau R, et al. Association study in a Sardinian sample between bipolar disorder

and the nuclear receptor REV-ERBalpha gene, a critical component of the circadian clock system. Bipolar Disord. 2009;11:215–220.
