## supplementary table 6 for "Studying cellular functions in bipolar disorder: Are there specific predictors of lithium response?"

**Supplementary table 6: Reagents and antibodies**

1. Anti-Nestin (Life technologies, Cat # A24354)
2. Anti-Pax6 (Sigma-Aldrich, Cat # AB2237)
3. Anti-TOMM22, (Sigma-Aldrich, Cat # HPA003037)
4. B27 supplement without Vitamin A (Thermofisher-Gibco, Cat #12587-010)
5. Beta-mercaptoethanol (Thermofisher-Gibco, Cat # 21985-023)
6. BFGF (Thermofisher-Gibco, Cat # PHG6015)
7. Carbonyl cyanide m-chorophenyl hydrazone (CCCP) (Sigma-Aldrich, Cat # C2759)
8. Click-it Edu Alexa Fluor 488 imaging kit (Thermofisher-Invitrogen, Cat # C10337)
9. DAPI (40, 6-diamidino-2-phenylindole) (Thermofisher-Life Technologies, Cat # R37606).
10. DMEM/F12 (Thermofisher-Gibco, Cat #10565-018)
11. DNA isolation kit (Macherey-Nagel, Cat # 740951.50)
12. Fetal bovine serum (Thermofisher-Gibco, Cat # 10270106)
13. Glutamax (Thermofisher-Gibco, Cat # 35050-061)
14. Heparin (Sigma-Aldrich, Cat #H3149)
15. Knockout DMEM (Thermofisher-Gibco, Cat # 10829-018)
16. KOSR (Thermofisher-Gibco, Cat # 10828-028)
17. Lithium Chloride (Sigma-Aldrich, Cat # L7026)
18. Lonza Mycoplasma detection kit (Lonza, Cat # LT07-318)
19. Matrigel (Corning, Cat. #354277)
20. MitoTracker™ Deep Red FM (Thermofisher-Invitrogen, Cat # M22426)
21. N2 supplement (Thermofisher-Gibco, Cat #17502-048)
22. Non-Essential Amino Acids (Thermofisher-Gibco, Cat # 11140-050)
23. Paraformaldehyde (Sigma-Aldrich, Cat # P6148)

24. Penicillin-Streptomycin (Thermofisher-Invitrogen, Cat # 15140-122)
25. Propidium Iodide dye (Thermofisher-Invitrogen, Cat # P3566)
26. Real Time PCR (q-PCR) system (Thermofisher, Cat # AB7500)
27. RNase A (Thermofisher-Invitrogen, Cat # 12091021)
28. RPMI-1640 (Himedia, Cat # AL060A)
29. Secondary antibody Alexa flour 488 donkey anti-mouse (Life Technologies, Cat # A24350)
30. Secondary antibody Alexa flour 594 donkey anti-rabbit (Life Technologies, Cat # A24343)
31. StemPro Accutase (Thermofisher-Gibco, Cat # A1110501)
32. SuperScript™ VILO™ cDNA Synthesis Kit (Thermofisher-Invitrogen, Cat # 11754050)
33. Sytox Green (Thermofisher-Invitrogen, Cat # S7020)
34. Taqman gene expression -housekeeping gene assays (Thermofisher-Applied Biosystems, Cat # 4448485)
35. Taqman gene expression master mix (Thermofisher-Applied Biosystems, Cat # 4369016)
36. Taqman gene expression -target gene Assays (Thermofisher-Applied Biosystems, Cat # 4331182)
37. Triton X-100 (Invitrogen, Cat # A24352)
38. Trizol (Thermofisher-Ambion, # 15596-026)
39. Valproic acid sodium salt (Sigma-Aldrich, Cat # P4543)
40. Vectashield (Vector labs, Cat # H-1000)
41. Verapamil (Sigma-Aldrich, Cat # V4629)
