## supplementary table 2 for "Studying cellular functions in bipolar disorder: Are there specific predictors of lithium response?"

### Supplementary table 2. Clinical and demographic characteristics of LCL study

| Variables | Total (N=25) | Responders (N=16) | Non-responders (N=9) | Chi square or z value | p value |
|---|---|---|---|---|---|
| Age at assessment | 40.2(12.8) | 37.6(10.5) | 44.8(15.6) | -1.0[a] | 0.3 |
| Gender-M: F (M %) | 15:10(60%) | 8:8(50%) | 7:2(77.8%) | 1.9[b] | 0.16 |
| Duration of illness (months) | 239.2(146.9) | 214.8(93.3) | 282.7(212.3) | -0.19[a] | 0.8 |
| Duration of lithium treatment (months) | 104.8(83.6) | 114.9(91.8) | 81.8(60.5) | -0.8[a] | 0.4 |
| Age of onset | 20.4(6.0) | 20.4(7.3) | 20.4(3.5) | -0.8[a] | 0.3 |
| Number of hospitalizations | 3.7(4.8) | 3.8(5.8) | 3.5(2.2) | -0.9[a] | 0.3 |
| Psychotic symptoms[#] | 20(83.3%) | 13(81.3%) | 7(87.5%) | 0.15[b] | 0.69 |
| Total no. of episodes | 8.4(6.2) | 7.6(5.0) | 10.1(8.4) | -0.7[a] | 0.4 |
| No. of manic episodes | 6.7(5.4) | 5.8(4.3) | 8.5(7.5) | -0.7[a] | 0.5 |
| No. of depression episodes | 1.8(2.5) | 1.5(2.2) | 2.5(3.2) | -0.7[a] | 0.4 |
| No of mixed episodes | 0.3(0.6) | 0.2(0.5) | 0.4(0.7) | -0.5[a] | 0.5 |
| Family H/o BD | 15(62.5%) | 8(50%) | 7(87.5%) | 3.5[b] | 0.06 |
| Family H/o psychosis | 3(13%) | 1(6.7%) | 2(25%) | N/A | N/A |
| Suicide attempt[#] | 2(9.1%) | 2(13.3%) | 0 | N/A | N/A |
| Onset episode | Mania- 19 (76%) | 11 (71.4%) | 8 (85.7%) | 0.56[b] | 0.45 |
| | Depression-6 (24%) | 5 (28.6%) | 1 (14.3%) | 0.56[b] | 0.45 |
| ALDA total score | 5.6(3.0) | 7.5(0.6) | 2.2(2.4) | -4.1[a] | 0.000* |
| A score | 7.88(3.1) | 9.8(0.5) | 4.4(2.9) | -4.4[a] | 0.000* |
| B score | 2.7(1.2) | 2.3(0.7) | 3.5(1.4) | -2.1[a] | 0.03* |

[#] Lifetime History, *p<0.05 (statistically significant); Values are mean (±SD), or n (%). [a] Mann-Whitney U test or [b] Pearson chi square were utilized to calculate p values across the variables between the responders and non-responders.
