## supplementary table 1 for "Studying cellular functions in bipolar disorder: Are there specific predictors of lithium response?"

**Supplementary table 1: Important studies related to mitochondrial function, cell death and cell proliferation in bipolar disorder**

| Author and year | Sample | Methodology | Significant results |
|---|---|---|---|
| **Studies related to mitochondrial function** | | | |
| Konradi et al., 2004 [1] | 9 BD and 10 control hippocampus from PM brains | Gene array to study mRNA expression in BD & control hippocampus | Expression of nuclear mRNA coding for mitochondrial proteins regulating oxidative phosphorylation in complexes I-V & ATP- dependent process were downregulated in hippocampus of BD. |
| Iwamoto, Kakiuchi, Bundo, Ikeda, & Kato, 2004 [2] | PM brain tissues:11 BD and 15 controls;<br>LCLs: 14BD & 11 controls | mRNA levels analysis in BD PM brain tissues and LCLS | Altered mRNA levels of proteins involved in aberration of protein translocation system into mitochondria (affecting mitochondrial function) in LCLs and brain tissues of BD. |
| Andreazza, Shao, Wang, & Young, 2010 [3] | 15 each post-mortem DLFC from BD patients & control | Investigated ETC complex I activity & oxidative damage to mitochondrial proteins along with levels of complex I subunit NDUFS7 | Levels of NDUFS7 & complex I activity decreased significantly in BD. Protein oxidation & 3-nitrosine increased in BD compared to controls. |
| Regenold et al., 2012 [4] | PM brain cortex tissue from 15BD and controls | Studied Hexokinase1 (HK1) attachment to outer mitochondrial membrane (OMM) in BD brain tissue | Decreased HK1 attachment in BD compared to controls. HK1 attachment to OMM, a critical feature of brain energy metabolism and survival of neurons through prevention apoptosis. |
| Gubert et al., 2013 [5] | Plasma and PBMC samples from 12BD and 30HC. | Evaluated oxidative stress marker in plasma and ETC complex activities in PBMCs of BD | No significant difference in oxidative stress markers and ETC complex activities between BD and HC. |
| de Sousa et al., 2015 [6] | 24 HC and 25 BD; Patients treated with lithium for 6 weeks | Evaluated leukocyte ETC complex activities in BD & effect of lithium on ETC | No significant differences in mitochondrial ETC complex activities between BD and HC. Lithium treatment significantly increased mitochondrial complex I activities. |
| Yoshimi et al., 2016 [7] | CSF: 54 BD & 40 controls;<br>PM brain tissue: 35BD & 34HC. | Evaluated the association of iso-citrate with BD | Iso-citrate levels significantly Increased in CSF from BD. mRNA levels of iso-citrate dehydrogenase in BD PM tissue was low compared to controls |

| | | | |
|---|---|---|---|
| Scaini et al., 2017 [8] | PBMC from 16BD and 16HC | Analysed levels of mRNA, protein and activity of mitochondrial related factors in PBMCs of BD and HC. | Levels of anti-apoptotic proteins and citrate synthase activity were significantly lower, while caspase activity was higher in PBMC of BD.<br>Levels of mRNA, protein related to mitochondria fusion were lower & related to fission were higher in PBMC of BD.<br>Showed mitochondrial dynamics & cell death pathway activation in BD, supporting link between mitochondria & pathophysiology of BD**.** |
| Bosetti et al., 2002 [9] | Rat-Lithium treatment for 7 days (acute) or 42 days (chronic). | Gene expression analysis of lithium treated rat brain | Chronic treatment at therapeutic concentration altered expression of several genes regulating mitochondrial enzymes in rat brain. |
| Lai, Zhao, Warsh, & Li, 2006 [10] | Human SH-SY5Y neuroblastoma (1 or 7 days of treatment) | Looked into the effect of lithium (1mM) or valproate (0.6mM) in stress induced human neuroblastoma cells | Pretreatment of SH-SY5Y cells for 7 days with lithium or valproate significantly reduced rotenone or $H_2O_2$ induced cytochrome C release, caspase activity & cytotoxicity and upregulated BCL2 protein level.<br>No effect was reported on 1 day treatment with the lithium or valproate. |
| Washizuka, Iwamoto, Kakiuchi, Bundo, & Kato, 2009 [11] | 1) LCLs from 25 BDI, 10 BDII & 33 HC.<br>2) 4 HC LCLs for lithium (0.75mM) or VPA (100ug/mL) treatment experiment | Studied gene expression of NDUFV2 in LCLCs of BD & HC; and after treatment of HC LCLS with lithium or VPA for 24hrs or 7 days | 1) NDUFV2 gene expression was significantly downregulated in BDI and upregulated in BDII compared to HC.<br>2) VPA treatment significantly increased NDUFV2 compared to vehicle. |
| Maurer, Schippel, & Volz, 2009 [12] | Human PM brain cortex from 5 Controls treated with lithium (0.1mM-10mM) for 10min | Investigated the effect of lithium on respiratory chain enzyme activities after exposure to lithium in human brain tissue | ETC complexes were significantly increased dose dependently by lithium with max at 1mm.<br>Succinate dehydrogenase was significantly increased at higher concentration of lithium |

| | | | |
|---|---|---|---|
| Bachmann et al., 2009 [13] | 1) SH-SY5Y cell line;<br>2) Brains from adult male Wistar Kyoto rats | Examined the effects of mood stabilizers on mitochondrial function and against mitochondrial mediated neurotoxicity | Cell respiration rate was enhanced by long term treatment with lithium or VPA. Mitochondrial function (membrane potential & oxidation) was enhanced by chronic lithium or VPA treatment in SH-SY5Y cells. *In vivo*: long-term treatment with lithium or VPA at therapeutic concentration prevented methamphetamine (meth) induced toxicity at the mitochondrial level (mitochondrial cytochrome c, anti-apoptotic Bcl-2/Bax ratio and COX activity).<br>Oligo array analysis: pre-treatment with lithium or VPA prevented meth induced dysregulation of gene expression of proteins related to apoptotic pathway and mitochondrial functions. *BCL2* expression increased after 6 days treatment. |
| Cataldo et al., 2010 [14] | 1) PM prefrontal cortex: 10 BD & 10 controls;<br>2) Fibroblasts: 8 BD & 8 HC;<br>3) LCLs: 6 BD & 6 HC | Evaluated structure and distribution of mitochondria in BD compared to controls; Effect of therapeutic dosage of lithium in fibroblast after 5 days treatment | Ultra-structure examination revealed smaller mitochondrial areas in BD brain PFC.<br>Altered mitochondria morphology & distribution was reported for BD fibroblasts and LCLs. Significant differences in cytochrome C distribution in BD fibroblasts.<br>No significant differences in mitochondrial differences in either groups on treatment. However, significant difference for distribution of mitochondria between treated BD and treated HC. |
| Sitarz et al., 2014 [15] | Fibroblasts from 5 POLG-deficient patients & 3 HC | Effect of 10mM VPA for 10 days on mitochondria associated proteins | VPA treatment increased mtDNA copy number. Protein levels of genes involved in mtDNA maintenance (POLG), mitochondria biogenesis & OXPHOS (COX2) increased significantly. |
| da Costa, Kormann, Galina, & Rehen, 2015 [16] | Neural progenitor cells from human embryonic stem cells | Effect of VPA (0.01,0.1, 1mM) for 24hours on cell size, mitochondrial morphology and function | Cell size and mitochondrial morphology changes after 1mM VPA treatment. Mitochondrial membrane potential (MMP) decreased with 1mM VPA. |

| | | | |
|---|---|---|---|
| Mertens et al., 2015 [17] | 6BD and 4HC dentate gyrus neurons derived from fibroblasts | Studied the mitochondria morphology and function in derived neurons from BD and HC. | Increased MMP and mitochondrial gene expression in BD neurons.<br>Size of neuronal mitochondria was smaller in BD compared to HC.<br>Lithium treatment increased mitochondrial size in lithium responsive neurons, whereas MMP remained unaffected. |
| Kakiuchi et al., 2005 [18] | 30 controls and 27 BD frontal cortex from PM brain tissue | Examined mitochondrial DNA (mtDNA) copy number in BD | No significant difference in mtDNA copy number of PM brain tissues between BD and controls. |
| Vawter et al., 2006 [19] | PM brain tissues from 9BD & 20controls | Analysed mitochondrial related gene expression and mtDNA copy numbers in BD and controls | 1) Mitochondrial gene & nDNA encoded mitochondrial genes were differentially expressed in BD. Mitochondrial related gene expression different in BD with lithium prescription Vs. BD without lithium at the time of death. The mtDNA copy number was non-significantly increased in BD compared to controls. |
| Sabunciyan et al., 2007 [20] | PM frontal cortex from 40BD & 44 controls | Examined mtDNA copy number in BD & controls | No significant difference in mtDNA copy number of PM brain tissues between BD and controls. |
| Torrell et al., 2013 [21] | PM brain tissues from 15BD & 15 controls | Examined mtDNA copy number and MT-ND1 gene expression in BD & controls | MT-ND1 gene expression was significantly increased in BD compared to controls. No significant difference for mtDNA content in PM brain tissues between BD and controls |
| C. C. Chang, Jou, Lin, & Liu, 2014) [22] | Leukocyte from 40 BD & 70 HC | Investigated mtDNA & oxidative damage in BD and HC leukocytes. | Leukocyte mtDNA copy number in BD was significantly lower than HC. Mitochondrial oxidative damage was significantly higher in BD compared to controls. |
| de Sousa et al., 2014) [23] | Leukocyte from 24 HC & 23 BD in depressive episode. | Evaluated mtDNA content in BD & HC. And tested if the content in BD varied after 6weeks of lithium treatment. | No significant difference in mtDNA copy number between BD and HC at baseline. No difference was reported even after 6 weeks of lithium treatment in BD cases. |
| Gabriel R. Fries et al., 2017 [24] | Peripheral blood samples from 22 BDI & 20 HC | Evaluated mtDNA copy number in BD & HC | The mtDNA copy number distribution in peripheral blood was significantly different in BD compared to HC. (Notably high variability in distribution was reported for BD group). |

| | | | |
|---|---|---|---|
| David Stacey et al., 2018 [25] | Peripheral blood samples from 50 BD (10 lithium responders and 40 lithium non-responders) | Comparison of mRNA expression levels between lithium responders and non-responders | 43 mRNA levels downregulated in lithium responders- includes mitochondrial encoded genes- MT-ND1, MT-ATP6, MT-CyB.<br>Genes involved in mitochondrial function pathway (ETC, OXPHOS) overexpressed. |
| **Studies related to cell death** | | | |
| Shao & Vawter, 2008 [26] | PM DLPFC from 29 BD and 27 controls | Gene expression array analysis in brain tissues from BD & HC | Genes involved in nervous system development, cell growth, & cell death were dysregulated. |
| McCurdy et al., 2006 [27] | Olfactory mucosa from 8 BD & 10 HC | Examined the rate of cell death in BD compared to HC | Cell death was significantly more in BD compared to HC. |
| F M Benes, Matzilevich, Burke, & Walsh, 2006 [28] | PM brain tissues from 9 BD& 10 controls. | Gene expression array analysis in brain tissues from BD and controls | 19 of 44 genes related to apoptosis were upregulated. Antioxidant related genes were downregulated. |
| Herberth et al., 2011 [29] | PBMCs from 16 BDI, 16BDII & 32HC; Validation in 7 BDI, 7 BDII & 14 HC. | Tested proteome profiling in PBMCs from BD & HCs. And effect of BD serum analytes on PBMCs from HC. | Proteome profiling of PBMC revealed differentially expressed proteins involved in cell death and survival pathways.<br>Addition of BD serum analytes on PBMCs from HC subjects reduced cell survivality. |
| Kazuno et al., 2013 [30] | 1) LCLs from monozygotic twins discordant for BD,<br>2) 8 LCLs each from BD & HC to validate | Evaluated protein markers in whole cell lysate derived from LCLs of twins & validated. | Several proteins involved in cell death & glycolysis was significantly differentially expressed between the patient and the co-twin. Case- control analysis validated only upregulation of PGAM1 (involved in energy metabolism) in BD cases. |
| Gabriel Rodrigo Fries et al., 2014 [31] | PBMC from 10 BD and 7 HC | Assessed cell death & viability in PBMCs of BD & HC | Cells in early apoptosis was significantly higher in BD.<br>No significant difference in cell viability, late apoptosis & necrosis between BD and HC. |
| Marianthi, Olga, Aristotelis, Nikolaos, & Fragiskos, 2015 [32] | Skin fibroblasts from 10 BD & 5 HC | Transcriptome profiling in fibroblasts from BD and HC | Genes involved in positive regulation of apoptotic process & cell cycle were differentially expressed in BD compared to HC. |

| | | | |
|---|---|---|---|
| Wollenhaupt-Aguiar et al., 2016 [33] | Neurons differentiated from neuroblastoma cell line (SH-SY5Y) challenged with serum of 12 BD or 6 HC | Investigated whether biochemical changes in the serum of patients induces neurotoxicity in neuronal cell cultures. | Reduced neurite density in neurons treated with serum of BD patients. Neurons challenged with serum of late stage patients showed significant decrease in cell viability |
| Xiaohua Li, Bijur, & Jope, 2002 [34] | Review article | Reviewed articles related to lithium or VPA treatment in human cell lines | Higher concentration of lithium reduced GSK3B activity & blocked facilitation of apoptosis.<br>VPA provided protection from apoptosis by inhibiting pro-apoptotic (such as reduced caspase 3 activity). |
| A. J. Kim, Shi, Austin, & Werstuck, 2005 [35] | Human hepatocarcinoma cell line (HEPG2) | Studied the ER stress induced dysfunction after pre-treatment with 0.5mM VPA for 18hours | Pre-treatment with VPA increased the cellular resistance to ER stress induced dysfunction and protects from apoptosis by inhibiting GSK3B. |
| Lai et al., 2006 [10] | Human SH-SY5Y neuroblastoma (1 or 7 days of treatment) | Looked into the effect of lithium (1mM) or valproate (0.6mM) in stress induced human neuroblastoma cells | Pretreatment of SH-SY5Y cells for 7 days with lithium or valproate significantly reduced rotenone or $H_2O_2$ induced cytochrome C release, caspase activity& cytotoxicity and upregulated BCL2 protein level. No effect was reported on 1 day treatment with the lithium or valproate. |
| Wilot et al., 2007 [36] | Hippocampal slices of rats | Evaluated neuroprotective effect of lithium & VPA against ATP induced cell death in rat hippocampus | ATP induced cell death was significantly reduced by lithium or VPA treatment at therapeutic dosage in both *in vitro* (acute) and *in vivo* (chronic) experiments. |
| Go et al., 2011 [37] | Neuronal progenitor cells (NPC) from embryonic brain of rats | Examined regulation of apoptotic cell death in rat NPCs by VPA | VPA (0.2, 0.5mM) treatment decreased NPC cell death after growth factor withdrawal or H2O2 stimulated peroxide conditions. VPA upregulated BCL-XL protein & mRNA levels in concentration dependent manner and suppressed Bax levels. The result was confirmed by *in vivo* in developing rat brains. |

| | | | |
|---|---|---|---|
| Lowthert et al., 2012 [38] | 8 weeks of open label lithium study in 20 BD patients | To assess the change in gene expression in peripheral blood of lithium responders and non-responders | 127 genes differentially expressed between responders and non-responders.<br>Pathway analysis showed regulation of apoptosis was significantly affected.<br>Upregulation of anti-apoptotic gene *BCL2* and downregulation of pro-apoptotic genes (*BAD*, *BAK1*) in responders and inverse relation in non-responders after 4 weeks. |
| Gawlik-Kotelnicka, Mielicki, Rabe-Jabłońska, Lazarek, & Strzelecki, 2016 [39] | Human neuroblastoma cell line (SH-SY5Y) | Assessed the effect of lithium (0.5 & 0.7mmol/L) for 24 hours in neuroblastoma cells. | Cell viability was significantly higher in therapeutic treated lithium samples than vehicle. |
| Del Grosso et al., 2016 [40] | Human oligodendrocyte cell line | Effect of lithium pre-treatment on cell viability | Psychosine induced autophagy in cells was rescued on lithium pre-treatment, by increasing cell viability. |
| Z. Li et al., 2017 [41] | Human neuroblastoma cell line (SH-SY5Y) | Effect of VPA in ER stress induced neuroblastoma cell lines on exposure to thapsigargin (TG) and on neuroprotection. | VPA treatment improves cell viability and reduces cell apoptosis in cell exposed to TG. ER stress induced apoptosis response protein were inhibited by VPA treatment. VPA upregulated the ratio of BCL2/Bax proteins in SH-SY5Y cells. VPA promotes cell proliferation through PI3K, AKT, GSK3B pathways. |
| Breen et al., 2016 [42] | LCLs from 23 Caucasian individual (8 BD lithium responders, 8 BD lithium nonresponders, 7 HC) | Exploring the effect of lithium 1mM (7 days) on transcriptome levels in LCLS from BD lithium response patients | Differential gene expression in apoptosis signalling system, defence response, protein processing pathways and response to ER stress pathways were discovered on treatment with lithium. |
| **Studies related to cell proliferation** | | | |
| McCurdy et al., 2006 [27] | Biopsies of olfactory mucosa from 8 BD & 10 HC | Explored the cell proliferation rates in BD compared to HC | No significant differences in mitosis between the BD and HC, however 11 genes involved in cell proliferation & 4 in neurogenesis were differentially expressed in BD. Cell death was significantly more in BD compared to HC. |

| | | | |
|---|---|---|---|
| F. M. Benes et al., 2007 [43] | Hippocampus tissues from PM brain of 7 BD & 7 controls | Gene expression profiling of brain tissues from BD and controls | *GAD67* (glutamate decarboxylase 67) gene expression was significantly decreased in BD than controls and *CCND2* (Cyclin D2) is known to regulate GAD67. *CCND2* was significantly downregulated in CA2/3 region of brain tissues in BD. |
| Francine M Benes, Lim, & Subburaju, 2009 [44] | Hippocampus tissues from PM brain of 7 BD & 7 controls | Evaluate the expression profiling of genes involved in G1 & G2 checkpoints of BD | Genes associated with transcriptional complex & G1 or G2 checkpoint of cell cycle regulation in BD was differentially expressed. Gene included *CCND2*, *CDK9* for G1 check point & *P53*, *CHK2*, *CCNE* for G2 checkpoint. |
| Marianthi et al., 2015 [32] | Skin fibroblasts from 10 BD & 5 HC | Transcriptome profiling in fibroblasts from BD and HC | Genes involved in positive regulation of apoptotic process & mitotic cell cycle were differentially expressed in BD compared to HC. |
| K. H. Kim et al., 2015 [45] | NPCs and matured neurons from 8 BD & 4 unaffected siblings | Transcriptomic microarray profiling in NPCs and matured neurons (early & late neurons) | Genes related to cell cycle were differentially expressed in BD late neurons compared to unaffected individuals. |
| Breen et al., 2016 [42] | LCLs from 23 Caucasian individual (8 BD lithium responders, 8 BD lithium nonresponders, 7 HC) | Exploring the effect of lithium 1mM (7 days) on transcriptome levels in LCLS from BD lithium response patients | Gene markers related to cell cycle and nucleotide excision repair were found to be differential in response to lithium between BD lithium responders and non-responders. |
| Mao, Hoang, & Dicorleto, 2001 [46] | Bovine aortic endothelial cells | Investigated the effect of lithium (5 & 10 mM) on regulation of cell cycle in bovine cells. | Lithium treatment increased G2/M cells without affecting cell viability up to 3 days whereas reduced thereafter.<br>Lithium increased mRNA and protein levels of p21, cyclin dependent kinase inhibitors.<br>Cyclin D mRNA expression was biphasic on lithium treatment--at 4 & 8 hours: It upregulated whereas at 24 & 48 hours: It was downregulated. |

| | | | |
|---|---|---|---|
| Sun et al., 2007 [47] | Human prostate cancer cells | Studied the effect of lithium in cell proliferation of prostate cancer cells. | Lithium (10mM) significantly inhibited cell proliferation at 72 hours treatment. Lithium significantly increased the percentage of cells in S phase & decreased cellular DNA replication.<br>Lithium altered expression of gene regulating DNA replication & Cell cycle. Cyclin A, Cyclin E, *P21* was downregulated whereas Cyclin D was upregulated. Protein level of Cyclin D was also up. |
| Seelan, Khalyfa, Lakshmanan, Casanova, & Parthasarathy, 2008 [48] | Human neuronal cell line (SK-N-AS) | Microarray expression profiling in human neuronal cell line after lithium (1.5mM) treatment for 33 days | Gene related to neuronal survival, growth, apoptosis, cell cycle regulation were differentially expressed on treatment with lithium. |
| Zanni et al., 2015 [49] | NPCs from mice | Studied the effect of lithium (1mM or 3mM) in mice NPCs | Lithium attenuated the effect of irradiation exposure induced cell cycle arrest in G1 and G2 phase. |
| Rattanawarawipa, Pavasant, Osathanon, & Sukarawan, 2016 [50] | Stem cells from human exfoliated deciduous teeth | Evaluated effect of lithium on cell proliferation in the cells after 3 days and 7 days treatment | Lithium significantly reduced colony forming unit ability/ proliferation in dose dependent manner. Lithium increased percentage of cells in subG0 phase, whereas decreased the percentage of cells in G1 phase after 3 days & 7 days of treatment. |
| Laeng et al., 2004 [51] | Rat neural stem cells | RNA profiling and protein analysis of rat neural stem cell after VPA or lithium treatment | Cell cycle regulating genes – *CCND2* was 5-6 fold increased on treatment by VPA, confirmed by protein analysis.<br>Lithium treatment for 3 days increased *CCND2* levels. |
| Catalano et al., 2005 [52] | Human papillary thyroid carcinoma cell line | Tested the effect of VPA on cell cycle phases of human carcinoma cell line. | VPA increased subG1 population in time and dose (0.5-3mM) dependent manner. Growth arrest in G1 phase was increased by VPA (1 &3mM) treatment.<br>Gene expression of *P21* and Cyclin A was increased. G2/M cell population was decreased non-significantly by VPA treatment. |
| X. N. Li et al., 2005 [53] | Human medullo-blastoma and | Investigated the effect of VPA (1 & 2.7mmol/L) on cell cycle phases of cell lines | VPA treatment caused cell cycle arrest for medulloblastoma cell line on day 7, i.e., significantly increased percentage of cells in G0/G1 phase & decreased cells in G2/M phase. |

| | | | |
|---|---|---|---|
| | neuroectodermal tumour cell lines | | |
| Wu & Guo, 2008 [54] | Immortalized human endometrial stromal cells | Examined the effect of VPA on cell cycle phases of stromal cell lines | VPA (3mM) for 16 hours treatment increased percentage of cells in G0/G1 phase and decreased cells in S phase & G2/M phases. |
| Witt et al., 2013 [55] | Primary murine prostate cancer cells (PCA) and fibroblasts | Studied the effect of VPA on *CCND2* expression in murine cell lines | VPA treatment highly increased *CCND2* gene expression in PCA cell line, however no effect of VPA on *CCND2* expression was seen in murine fibroblast. |
| Claudia Morich, 2016 [56] | Tumour cell lines | Expression profiling after VPA treatment | *CCND2* gene expression was significantly increased after VPA treatment. |
| Pietruczuk, Lisowska, Grabowski, Landowski, & Witkowski, 2018 [57] | T cells from 18 BD & 10 HC | Evaluated proliferation capacity & susceptibility to apoptosis in T cells and effect of lithium or valproate on these parameters | Cell cycle longer in BD compared to HC; reduced proliferation in lithium treated BD patients compare to HC and BD treated with VPA.<br>Cell cycle longer in patients treated with VPA compared to lithium treated patients.<br>*In vitro* exposure to VPA reduced cell division and cell proliferation irrespective of the disease state; lithium has no effect on proliferating capacity of T cells from BD patients. Higher doses of lithium shortened cell cycle.<br>Apoptosis higher in BD cells, lithium and VPA prevents apoptosis in T cells from BD.<br>BCl2 level: No significant difference between BD and HC |

3. Andreazza AC, Shao L, Wang J-F, Young LT. Mitochondrial Complex I Activity and Oxidative Damage to Mitochondrial Proteins in the

Prefrontal Cortex of Patients With Bipolar Disorder. Arch Gen Psychiatry. 2010;67:360.

15. Sitarz KS, Elliott HR, Karaman BS, Relton C, Chinnery PF, Horvath R. Valproic acid triggers increased mitochondrial biogenesis in

POLG-deficient fibroblasts. Mol Genet Metab. 2014;112:57–63.

41. Li Z, Wu F, Zhang X, Chai Y, Chen D, Yang Y, et al. Valproate Attenuates Endoplasmic Reticulum Stress-Induced Apoptosis in SH-

SY5Y Cells via the AKT/GSK3β Signaling Pathway. Int J Mol Sci. 2017;18.

56. Claudia Morich. The influence of valproic acid and the role of cyclin D2 in prostate cancer. Georg-August University, Göttingen, 2016.

57. Pietruczuk K, Lisowska KA, Grabowski K, Landowski J, Witkowski JM. Proliferation and apoptosis of T lymphocytes in patients with bipolar disorder. Sci Rep. 2018;8:3327.
